# The action of a cosmetic hair treatment on follicle function

**DOI:** 10.1101/290437

**Authors:** Graham A. Turner, Sarah E. Paterson, Fiona L. Baines, Andrew E. Mayes, David M. Reilly, Nicole M. Hudson, Tony Dadd, Amitabha Majumdar, Renu Kapoor, Jiayin Gu, Nitesh Bhalla, Fei Xu

**Author notes:** Correspondence: Dr Graham Turner, Unilever R&D Port Sunlight, Quarry Road East, Bebington, Merseyside, CH63 3JW, UK. Current affiliation: Hindustan Unilever Limited, Mumbai, India (Nitesh Bhalla). Co-author contact details: Sarah E. Paterson, Unilever R&D Port Sunlight, Quarry Road East, Bebington, Merseyside, CH63 3JW, UK; Fiona L. Baines, Unilever R&D Port Sunlight, Quarry Road East, Bebington, Merseyside, CH63 3JW, UK; Andrew E. Mayes, Unilever R&D, Colworth Science Park, Sharnbrook, MK44 1LQ, UK; David M. Reilly, Unilever R&D, Colworth Science Park, Sharnbrook, MK44 1LQ, UK; Nicole M. Hudson, Unilever R&D, Colworth Science Park, Sharnbrook, MK44 1LQ, UK; Tony Dadd, Unilever R&D, Colworth Science Park, Sharnbrook, MK44 1LQ, UK; Amitabha Majumdar, Unilever R&D Bangalore, 64 Main Road, Whitefield PO, Bangalore 560 066, India; Fax: N/A; Renu Kapoor, Unilever R&D Bangalore, 64 Main Road, Whitefield PO, Bangalore 560 066, India; Fax: N/A; Jiayin Gu, Unilever R&D Shanghai, 66 LinXin Road, Changning District, Shanghai, 200335, China; Fax: N/A; Nitesh Bhalla, Hindustan Unilever Limited, Mumbai HURC, B.D. Sawant Marg, Chakala, Andheri (E), Mumbai 400 099, India; Fax: N/A; Fei Xu, Unilever R&D Shanghai, 66 LinXin Road, Changning District, Shanghai, 200335, China; Fax: N/A.

## Abstract

**OBJECTIVE:** Human hair changes with age: fibre diameter and density decrease, hair growth slows and shedding increases. This series of controlled studies examined the effect on hair growth parameters of a new leave-on hair treatment (LOT) formulated with Dynagen^TM^ (containing hydrolysed yeast protein) and zinc salts.

**METHODS:** Hair growth data were collected from healthy women aged 18–65 years. The LOT’s effect on hair growth was measured in a randomized double-blind study and in hair samples; its effect on follicle-cell proliferation was assessed by quantifying Ki67 expression in scalp biopsies. The LOT’s effect on plucking force was determined in an *ex vivo* model. Dynagen’s effect on the expression of the tight-junction marker claudin-1 was analysed in cultured follicles. The effect on protease activity of zinc salts used in the LOT was examined *in vitro*.

**RESULTS:** Hair growth rate decreased with increasing subject age. The LOT significantly increased hair growth rate, fibre diameter, bundle cross-sectional area, Ki67 expression and the plucking force required to remove hair. Dynagen significantly increased claudin-1 expression in cultured follicles. Protease activity was reduced by zinc salts.

**CONCLUSION:** The Dynagen-based LOT increases hair-fibre diameter, strengthens the follicular root structure and increases hair growth rate.

## Introduction

Human hair grows in a cyclical, asynchronous pattern, exhibiting three key stages: anagen (growth), catagen (a transition phase) and telogen (resting phase). At the end of telogen the hair fibre is shed from the scalp in a process often referred to as exogen. As the hair follicle cycles, there is a natural amount of hair loss from the scalp, estimated to be 50–150 hairs over a 24-hour period [1].

A number of attributes of human hair are known to change as a result of ageing, including hair colour (greying), hair density (loss), hair diameter (reduced) and growth rate (slower) [2–10]. Alterations in hair-fibre properties with age have also been described, e.g. nanomechanical properties and tactile perception [11], as well as changes in the force required to pluck hairs [12]. Such changes may provide opportunities to ameliorate age-related changes in hair.

To date, attempts to mitigate the effects of age through measurable actions on the biology of hair have been mainly limited to the administration of pharmacological agents such as minoxidil and finasteride [13–15]. However, ingredients used in conventional cosmetic products may provide a means to support such medical interventions. From a biological perspective there are a number of approaches that could be undertaken to influence hair shedding rate and abundance. One would be to maintain the anagen phase, thus preventing the transition, via catagen, into telogen with the inevitable progression to the exogen (shedding) phase. For example, procyanidin B2 – a proanthocyanidin found in grape-seed extract and other plant materials – has been demonstrated to prevent the anagen to catagen transition by blocking transforming growth factor β2 signalling [16]. Once in telogen, the process of hair shedding commences immediately. Alternatively, as the breakdown of linkages between the hair fibre and the follicle structure is a proteolytic process [17], inhibition of protease activity during this phase may have the potential to reduce the rate of hair shedding. For example, a combination of Trichogen^®^ and climbazole has been shown to inhibit proteases found in the human hair follicle, and was associated with an increase in the force required to remove hair in an *ex vivo* skin model [18]. Trichogen includes zinc gluconate; zinc salts are known to inhibit protease activity [19].

This paper describes a series of studies that examined the effect of age on hair-growth parameters and the effect of a new bioactive incorporated into a leave-on treatment (LOT) formulated to target specific aspects of hair biology relating to hair loss and hair-fibre diameter, i.e. maintenance of the anagen phase and inhibition of proteases. The key active ingredient, Dynagen™ (Ashland Specialty Ingredients, Wayne, NJ, USA), contains yeast protein hydrolysate, which has been shown to increase the expression of the key structural hair-follicle proteins keratin 14 and collagen IV [20–24]. The bioactive formulation also contains zinc salts. The aim of the *in vitro*, *ex vivo* and clinical studies was to investigate the effects of the Dynagen plus zinc salts LOT on hair attributes related to hair loss, hair- fibre diameter and hair growth rate.

## Methods

### Hair growth

#### Effect of age on hair growth

Phototrichogram (PTG) scalp-hair measurements [25] were taken from 255 women (age range 18–65 years) who were in good general health, with no recognized or diagnosed scalp or hair conditions. Data were collected under standardized conditions at Alba Science Ltd (Edinburgh, UK) in accordance with the Declaration of Helsinki following receipt of informed consent and with ethical approval from the Reading Independent Ethics Committee. Relationships were examined between the subject’s age and a range of PTG endpoints including total and anagen hair count, hair width and length, and hair growth rate determined over 48 hours. Data were first analysed comparing older (55–65 years) and younger (18–25 years) age groups; further analyses were conducted in the whole cohort. Statistical analyses were conducted using JMP 11.0 (SAS Institute Inc., Cary, NC, USA).

#### Clinical study: effect of active LOT on hair growth *in vivo*

A double-blind, randomized, placebo-controlled, parallel-group clinical study was conducted in healthy women aged 23–50 years who had no history of medically diagnosed hair loss, had at least 80% of their scalp hair in anagen phase at screening (as determined by pluck trichogram) and had given written informed consent. The study was conducted at Alba Science Ltd with ethical approval from Reading Independent Ethics Committee and in accordance with the Declaration of Helsinki. Following a 3-week lead-in (conditioning) phase, subjects were randomly allocated (1:1) to either an active LOT (1.5% Dynagen, 0.018% zinc sulphate heptahydrate; active LOT) or placebo (LOT without Dynagen or zinc salts) using a stratified randomization scheme in which subjects were balanced for age and number of anagen hairs (as determined by pluck trichogram performed at screening). During a 16-week test phase, subjects applied either active LOT or placebo to their whole scalp once daily for five consecutive days followed by two consecutive days without application, for a total of 80 applications per subject. Briefly, subjects washed their hair according to their normal habits using a commercially available shampoo and conditioner (Dove Daily Care shampoo and conditioner; Unilever, Bebington, UK). The supplied LOT was applied immediately after the hair was washed and before hair drying, and remained on the scalp for at least 20 hours before the next hair wash. Hair growth was measured at baseline and every 4 weeks until the end of the study using the Canfield PTG technique [25]. Statistical analysis was performed using SAS for Windows (version 9.4; SAS Institute).

### Hair-follicle studies

#### Hair growth in isolated human hair follicles

The Philpott isolated-follicle model [26] was used to assess hair growth. Briefly, follicular units containing anagen hair follicles were microdissected from hair grafts from the occipital scalp of healthy male adults (aged 20–45 years), obtained as discarded cosmetic surgery waste (Pioneer Hair Care Centre, Bengaluru, India) with informed consent and following local ethical committee approval (Unilever Independent Ethics Committee; e-IEC Reference ETH2013_HDT_HB1b 01). The epidermis and upper dermis were removed from the subcutaneous fat layer and the hair follicles were then gently dissected from the subcutaneous layer using fine forceps. Anagen hair follicles, identified by their well-rounded bulb structure, were placed in 500 μL of culture medium (William’s E medium [Sigma Aldrich Co, Gillingham, Dorset, UK], supplemented with GlutaMax-1 [Life Technologies, Carlsbad, CA, USA], 10 ng mL^−1^ hydrocortisone and 100 units mL^−1^ penicillin) in a 24-well plate (two follicles per well), and either 15 µL of 1.5% Dynagen or control (William’s E medium without Dynagen) were added to each well. The follicles were cultured in a humidified incubator at 37°C and 5% CO_2_ in air, and allowed to grow for 9 days, with the culture medium being changed every 2–3 days (Dynagen/control was added again at each change of medium), before fixing with 4% buffered formalin.

Following isolation and culture of the hair follicles, hair growth was measured by capturing images on alternate days for 9 days using a stereomicroscope (Olympus SZ40) with photo-capturing software, and analysed using Image-Pro Plus software (Media Cybernetics, Rockville, MD, USA). Each experiment was conducted with at least four follicular units per subject and the results presented are means of three subjects. Data were analysed using analysis of variance (ANOVA) followed by Dunnett’s test for multiple comparisons. Results were considered to be significant at *P*<0.05.

#### Hair-follicle markers

##### Ki67

The effect of the active LOT on expression of Ki67 – a marker of cell proliferation – was analysed using immunohistochemical techniques in the hair follicles from 4-mm punch biopsies of scalp skin (10 from each treatment group), taken following unblinding at the end of the clinical study described above. . The biopsies were snap frozen in OCT medium and stored at –80°C until required. For sectioning, samples were placed in a cryostat (Thermo Shandon Cryotome FSE/FE; Thermo Fisher, Paisley, UK) and serially cryosectioned into 7-µm thick sections. Sections containing anagen hairs – identified by their well-rounded bulb structure – were mounted on SuperFrost^®^ microscope slides (Fisher Scientific, Loughborough, UK) and stored at –80**°**C for later use.

For immunolocalization of Ki67, sections were stained using a commercially available avidin–biotin system (VECTASTAIN Elite ABC peroxidase kit PK-6102; Vector Laboratories, Peterborough, UK) according to the kit manufacturer’s protocol. Briefly, slides were air dried and fixed using Zamboni’s fixative for 10 minutes. Non-specific binding was blocked by application of normal horse serum (PK-6102; Vector Laboratories) diluted in tris-buffered saline. Sections were incubated at room temperature for 1 hour with mouse anti-Ki67 primary antibody (VP-K452; Vector Laboratories; 1:100 in tris-buffered saline). Biotinylated horse anti-mouse secondary antibody (PK-6102; Vector Laboratories) was then applied and the slides were incubated with the avidin–biotin reagent. ImmPACT AMEC red peroxidase substrate solution (SK-4285; Vector Laboratories) was applied until the desired stain intensity developed. Slides were counterstained using filtered 1:10 Gill’s #3 hematoxylin (1:100; PRC/13/3; Pioneer Research Chemicals, Colchester, UK).

Positively stained cells in the hair follicle and hair bulb were counted using a graticule with a 10 × 10 square grid, at a magnification of ×40. The percent area of Ki67-positive cells per unit area in the bulb and in the follicle were quantified by Nuance 3.0.2 Multispectral Imaging System software (Caliper Lifesciences, PerkinElmer, Beaconsfield, UK). Statistical analysis of Ki67 data was performed using a simple mixed ANOVA model including a fixed effect for treatment and a random effect for subject.

##### Claudin-1

The effect of Dynagen on tight junctions was assessed by measuring the expression of the tight junction-associated protein claudin-1 in isolated hair follicles *in vitro*. Isolated follicles were fixed following the follicular hair-growth study described above. Twenty regions were analyzed in each of three follicles taken from each of three volunteers per group. The tissues were cross-sectioned (3–5 µm thickness) and the hair-follicle sections dewaxed followed by boiling in citrate buffer (pH 6) for antigen retrieval. Sections were mounted on slides (two sections per slide), then incubated with 1% (w/w) bovine serum albumin (Sigma Aldrich Co) to minimize non-specific binding, followed by overnight incubation at 4°C with rabbit anti-claudin-1 antibody (51-9000, 1:10 dilution; Cell Signaling Technology, Danvers, MA, USA). Finally, sections were incubated with an Alexa Fluor^®^ 488-labelled goat anti-rabbit secondary antibody (1:15 dilution; Life Technologies) for 30 minutes at room temperature. Digital images were captured using an Olympus BX50 microscope with U- CMAD3 adapter, and analysed using ImageJ software (National Institutes of Health, Bethesda, MD, USA).

#### Measurement of plucking force

Excised pig skin pieces (1 cm × 1 cm), as described above, were incubated in high-glucose Dulbecco’s Modified Eagle’s Medium and GlutaMax^TM^-I (Life Technologies), supplemented with 10% foetal bovine serum, 100 U mL^−1^ penicillin and 100 μg mL^−1^ streptomycin. Active LOT or control (water) was applied to the skin at 0.2 mL cm^−2^ (LOT peptides: 3 mg cm^−2^). After application, the skin samples were cultured at 37°C for 48 hours, with a second application of LOT/control after 24 hours. The extraction force required to remove hairs from the pig skin was then determined using a Zwick Z005 displacement controlled tensile testing machine as described previously [18]. Thirty individual hair fibres were plucked from each piece of pig skin and the extraction force recorded. All plucking-force data were expressed as means ± standard errors of the mean. Statistical significance was examined using the Fit Model platform (JMP 11.0; SAS Institute); results were considered significant at *P*<0.05.

#### Protease inhibition in vitro

Protease inhibition by the zinc salts (0.0001% zinc gluconate, 0.018% zinc sulphate heptahydrate) used in the active LOT was evaluated versus zinc salt-free control using a commercial assay kit (EnzChek^TM^ Protease Assay kit; Molecular Probes, Eugene, OR, USA), as reported previously [18]. Results are expressed as means ± standard errors of the mean (3–5 replicates/sample). Statistical analysis was by Student’s *t* test, with results considered significant at *P*<0.05.

### Hair-fibre diameter

For single-fibre and switch-diameter measurements, hair switches (2 g, 25 cm long, double-bleached, round, European brown hair) were washed with 14% (v/v) sodium laureth sulphate and left to dry overnight at 20°C and 50% relative humidity (RH).

#### Measurement of single-fibre diameter

Fifty numbered fibres were cut from the top of a single switch and crimped. The cross-sectional diameter of each fibre was measured with a laser-scanning micrometer (Model FDAS770; Dia-Stron, Andover, UK) at five locations along the fibre and four rotating angles (0°, 90°, 180° and 270°) at each location, using the Fibre Dimensional Analysis System (Dia-Stron, Andover, UK). Each fibre was soaked for 1 hour in 200 μL of active LOT and left to dry overnight at 20°C, 50% RH. The cross-sectional diameter of each fibre was then measured again using the same method. The average diameter for each fibre before and after treatment was calculated from the two sets of 20 data points collected. Statistical analysis was performed using a one-tailed Student’s *t* test for paired samples (before and after treatment); *P*<0.05 was considered to be significant.

#### Measurement of switch diameter

The Hair Matrix Index for each of five base-washed hair switches was measured at three locations (5, 10 and 15 cm from the top) using a cross-section trichometer (HairCheck^®^; Haircheck International Co., Miami, FL, USA), as described by Cohen [27]. Active LOT (200 µL) was applied to each location where the thickness was measured previously and the switches were left to dry overnight at 20°C, 50% RH. The Hair Matrix Index at each location was then measured again using the cross-section trichometer and the cross-sectional area calculated. Statistical analysis was performed using a one-tailed Student’s *t* test for paired samples (before and after treatment); *P*<0.05 was considered to be significant. As obtained from the suppliers, Dynagen contains glycerol (30%), which is a humectant that can form hydrogen bonds with water to retain moisture, thus swelling fibres and increasing diameter. Therefore the experiments were repeated, applying 200 µL glycerol (30%) in place of the active LOT to determine the extent of fibre diameter increase by glycerol alone.

## Results

### Hair growth

#### Effect of age on hair growth

Statistically significant differences were found between younger (n=48; median age 22 years; range 18–25 years) and older (n=40; median age 59 years; range 55–65 years) women for total hair count (*P*<0.0001), total anagen count (*P*<0.0001), total telogen count (*P*<0.05), actual growth rate (*P*<0.0001), cumulative hair width (*P*<0.0001) and cumulative anagen hair width (*P*<0.0001), although not for average hair width or average anagen hair width (*P*=0.3073 and *P*=0.6085, respectively); data not shown. For all endpoints, the younger subjects had higher values than the older subjects. Analysis of the relationships between each of the PTG endpoints and age showed that, as expected, many of the endpoints were highly correlated with each other (e.g. anagen count and total hair count). However, the associations with age were generally much weaker. The highest Pearson correlation coefficients with age were approximately –0.3: cumulative hair width (r=–0.33), cumulative anagen width (r=–0.31), total hair count (r=–0.30), actual growth rate (r=–0.30) and total anagen count (r=–0.29); data not shown. The correlation coefficients were weaker still for age and average hair width and average anagen hair width (r=–0.09 and r=–0.07, respectively; data not shown).

The relationship between hair growth rate and age was examined in more detail using linear modelling in the entire cohort of 255 subjects (mean age 36.6 years, median age 33 years; age range 18–65 years), which showed that hair growth rate decreased with increasing age, by an average of 0.043 μm h^−1^ per year (Fig. 1).

**Figure 1.**
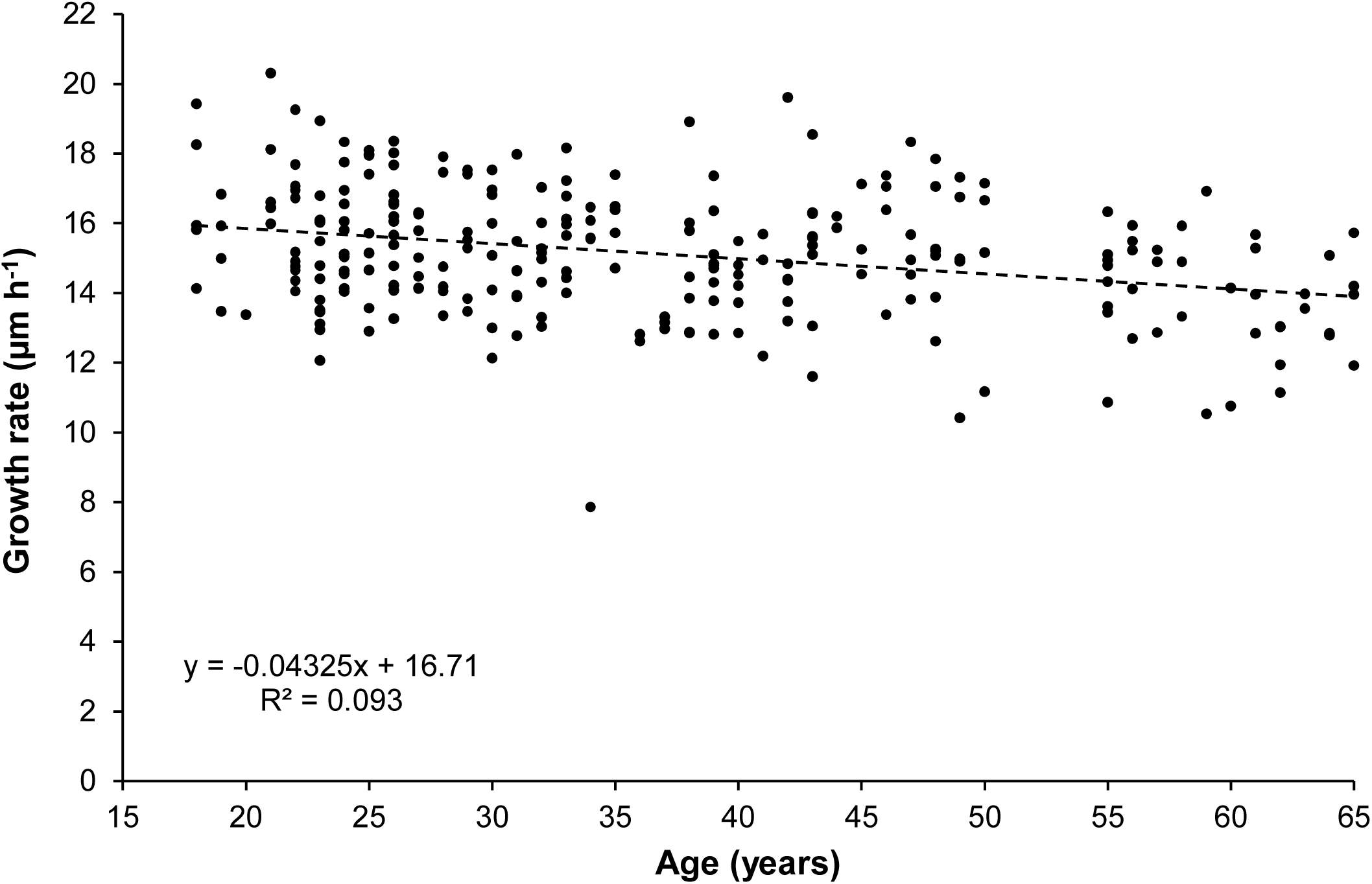
Effect of age on hair growth rate. Growth rate for each subject was determined over a nominal 48 hours, based on the actual elapsed time between individual measurements.

#### Effect of active LOT on hair growth *in vivo*

A total of 132 subjects completed the study; 70 in the control group and 62 in the active LOT treatment group (Fig. 2). In subjects treated with active LOT, hair growth rate over the 16-week study period was significantly greater than in subjects who used the control LOT (15.56 µm h^−1^ [95% confidence interval (CI) 15.31 to 15.81 µm h–1] vs 15.03 µm h^−1^ [95% CI 14.79 to 15.28 µm h^−1^]; *P*=0.0035; main effects model with least-squares mean estimates and baseline as covariate) (Fig. 3). There was no evidence that the difference between the treatments varied with time (*P*=0.601; data not shown). There were very few treatment-related adverse events reported in the study (n=48), all of which were minor in nature (e.g. irritation, pruritus); 46 were mild in severity and two were moderate.

**Figure 2.**
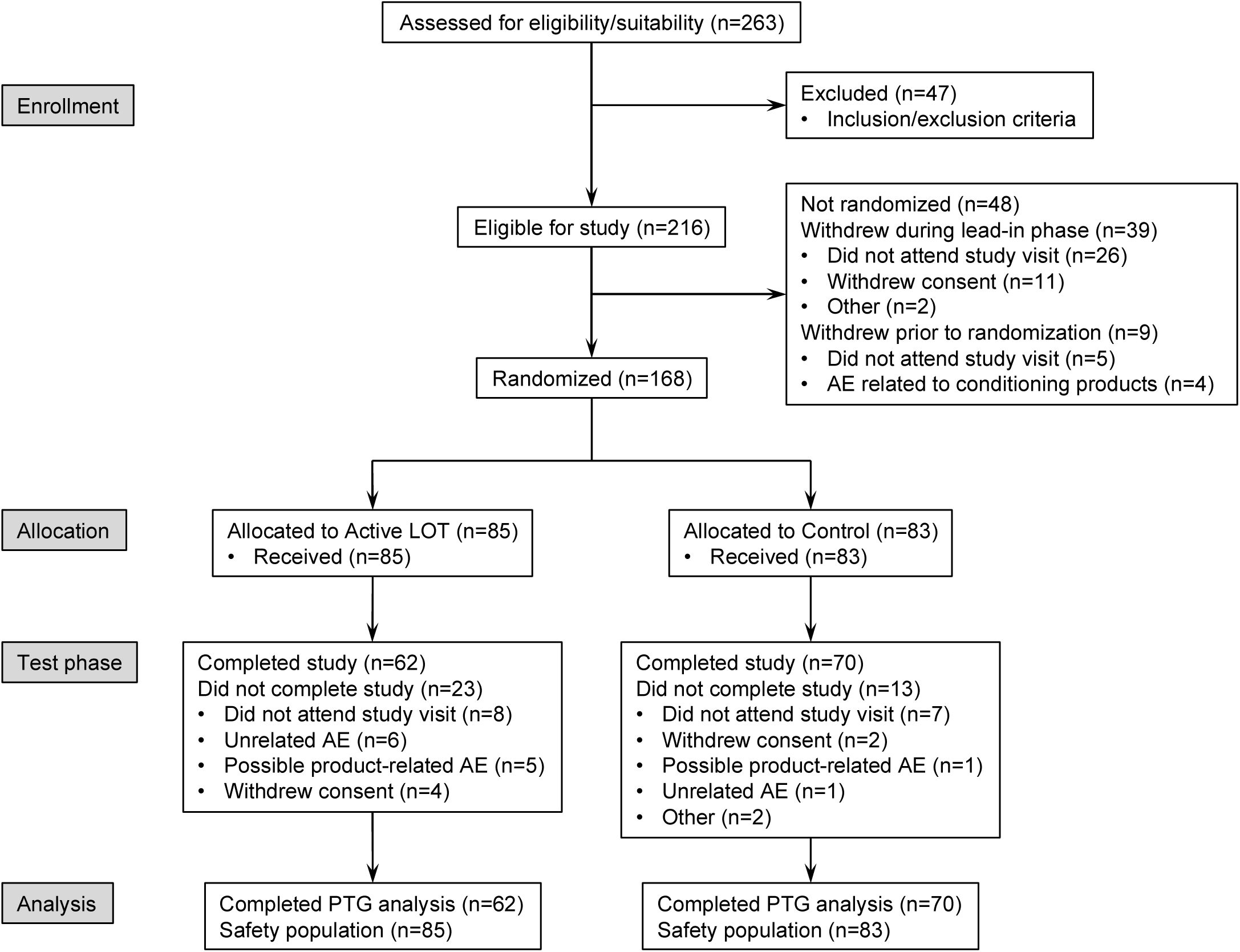
Flow diagram showing subject disposition in the clinical study. The main reasons for withdrawal of consent during the study were ‘could not attend further study visits’ (n=11) and ‘did not wish to comply with study restrictions’ (n=4). AE = adverse event; LOT = leave-on treatment; PTG = phototrichogram.

**Figure 3.**
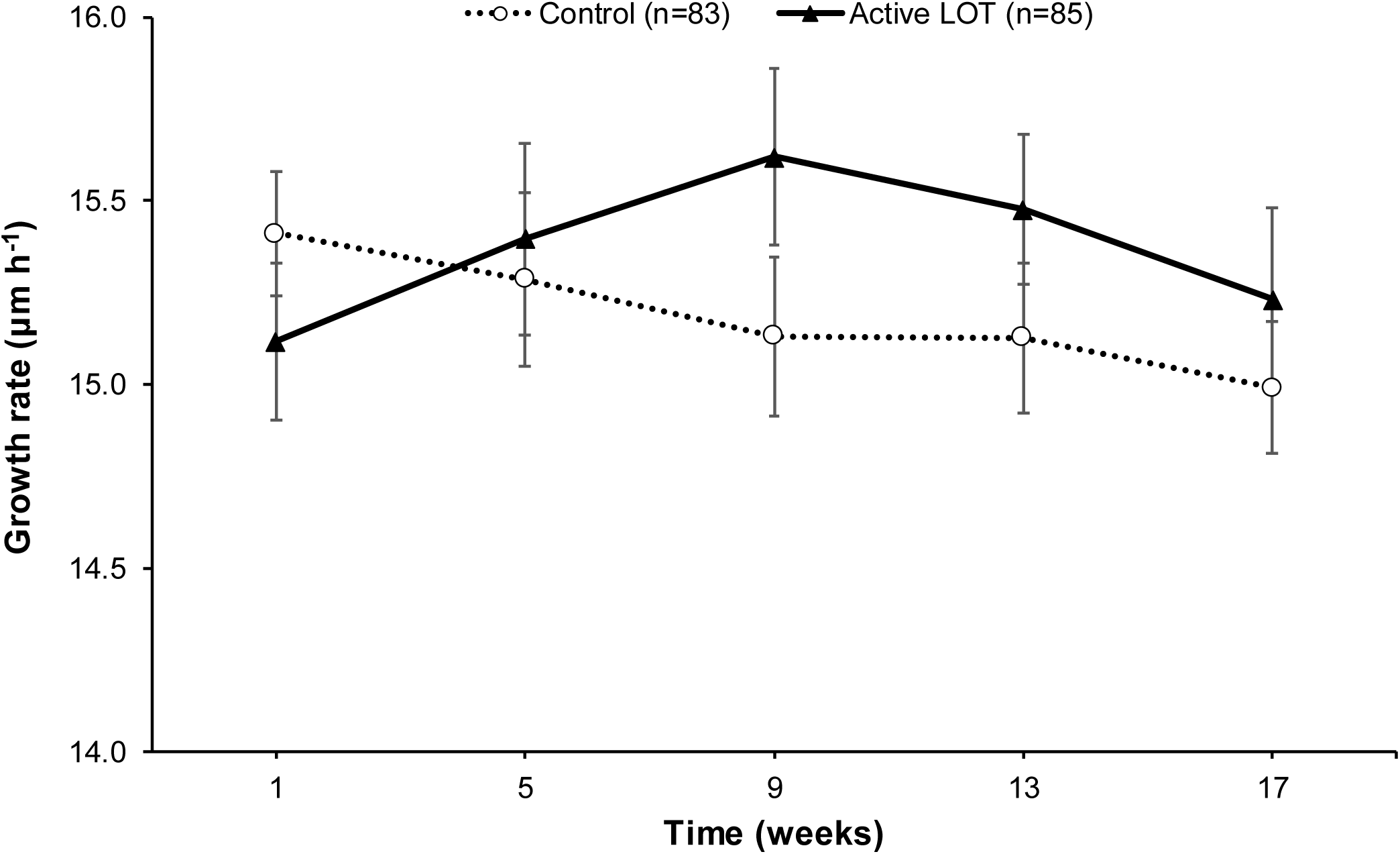
Hair growth rate over time in response to active and control LOTs. Results presented are means and standard errors of the mean. LOT = leave-on treatment.

### Hair follicle-related studies

#### Hair growth in isolated human hair-follicle model

Hair follicles cultured *in vitro* over a period of 9 days grew at a faster rate when treated with active Dynagen versus control (*P*<0.05; Fig. 4).

**Figure 4.**
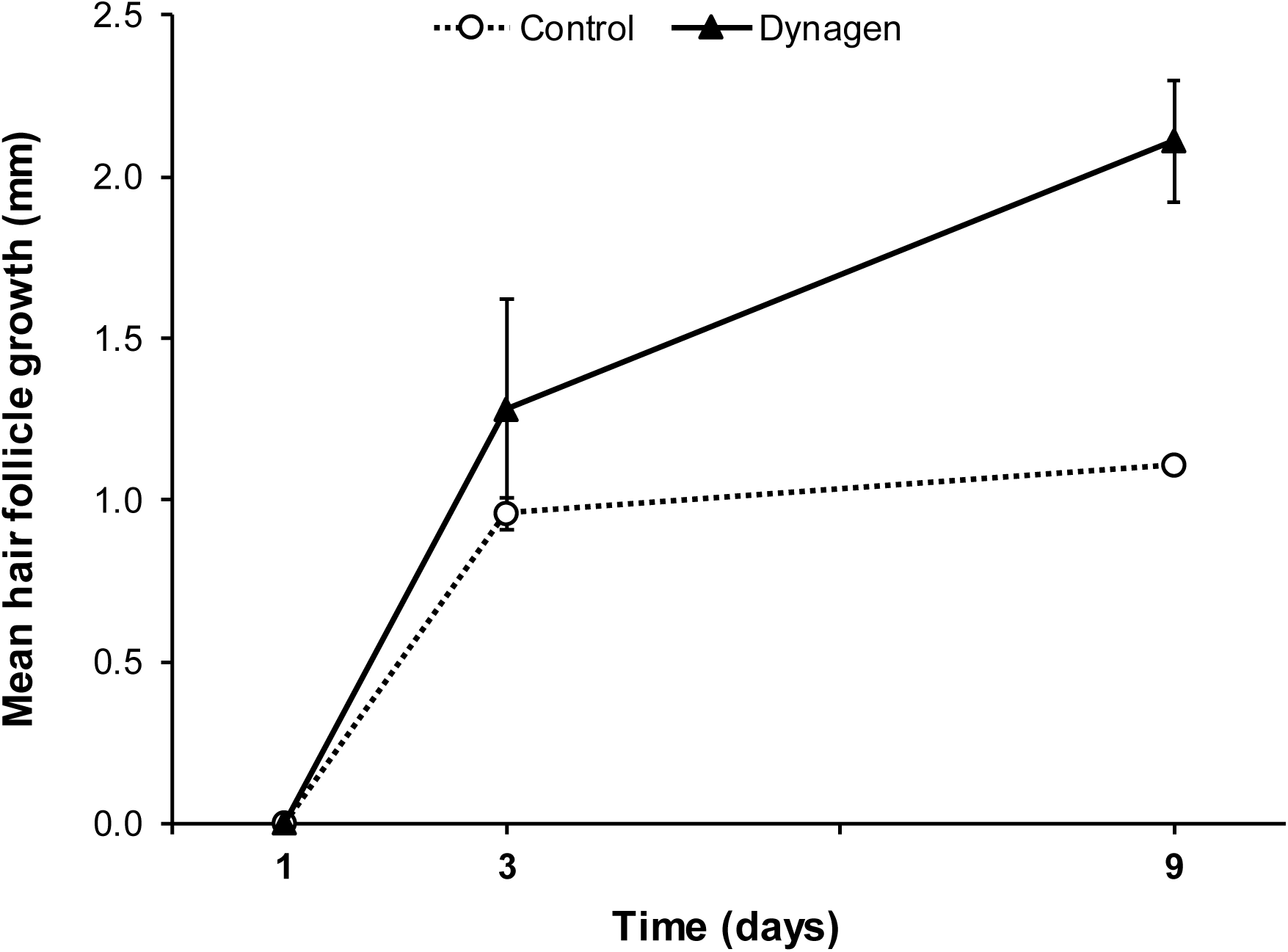
Effect of 1.5% Dynagen on the growth rate of cultured human hair follicles *in vitro*. Results presented are means and standard deviations of the means; data are from at least four follicular units per volunteer from three volunteers per group.

#### Follicle markers

##### Ki67 expression

There were no significant differences in mean visual count of the number of Ki67-positive cells in the bulb or follicle of hairs from subjects treated with active LOT compared with those receiving control LOT (mean cell counts 4.23 vs 3.91 in bulb [*P*=0.38] and 1.50 vs 1.37 [*P*=0.46] in follicle, respectively) (Fig. 5a). There was no significant between-group difference in the area of Ki67 expression in the follicle (active LOT 26.8% vs control LOT 25.5%; *P*=0.81) (Fig 5b); however, the expression of Ki67 measured in the hair bulbs from subjects who had been treated with the active LOT was statistically significantly greater than in those from subjects who had used the control LOT (*P*=0.032). Comparing the percent area of Ki67-positive cells in hair bulbs from active LOT-treated subjects with that in the control group showed that the difference in area of staining was 12% (64.0% vs 52.4%, respectively; estimated mean difference [control minus test] = –11.9; 95% CI: –1.1 to –22.5) (Fig 5b).

**Figure 5.**
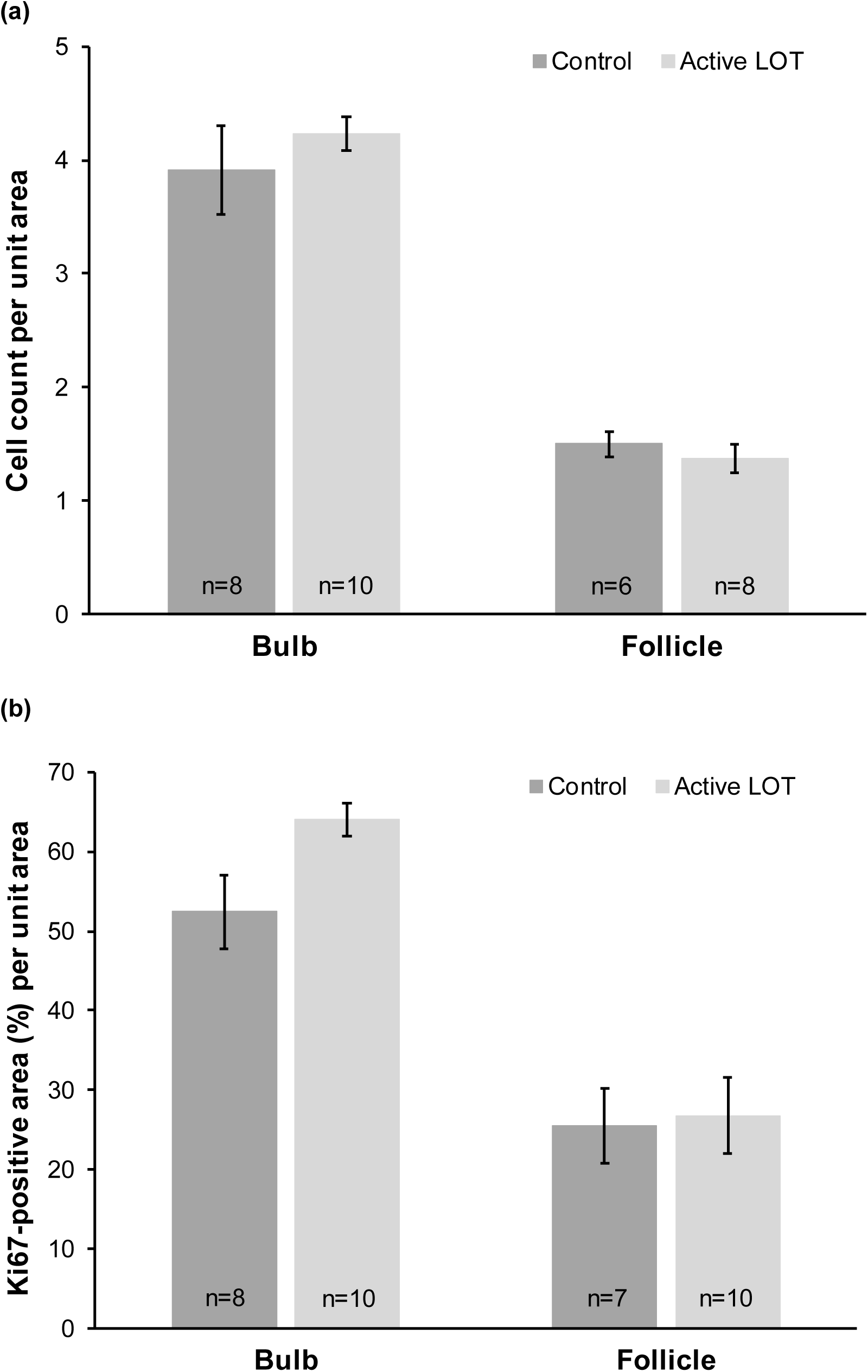
Extent of Ki67 expression in bulbs and follicles of hair from subjects receiving active or control LOTs. (a) Mean (± standard error of the mean) visual count of Ki67- positive cells per unit area. (b) Mean (± standard error of the mean) percent area of Ki67-positive cells per unit area. LOT = leave-on treatment.

##### Claudin-1 expression

Compared with control, Dynagen treatment was associated with increased expression of claudin-1 protein in the cultured hair follicles (Fig. 6a). Quantitative analysis with ImageJ using the cross-section of the hair follicle as the region of interest (Fig. 6b) also showed increased expression of claudin-1 in Dynagen-treated follicles (*P*<0.0001; ANOVA followed by Dunnett’s method), supporting the qualitative results.

**Figure 6.**
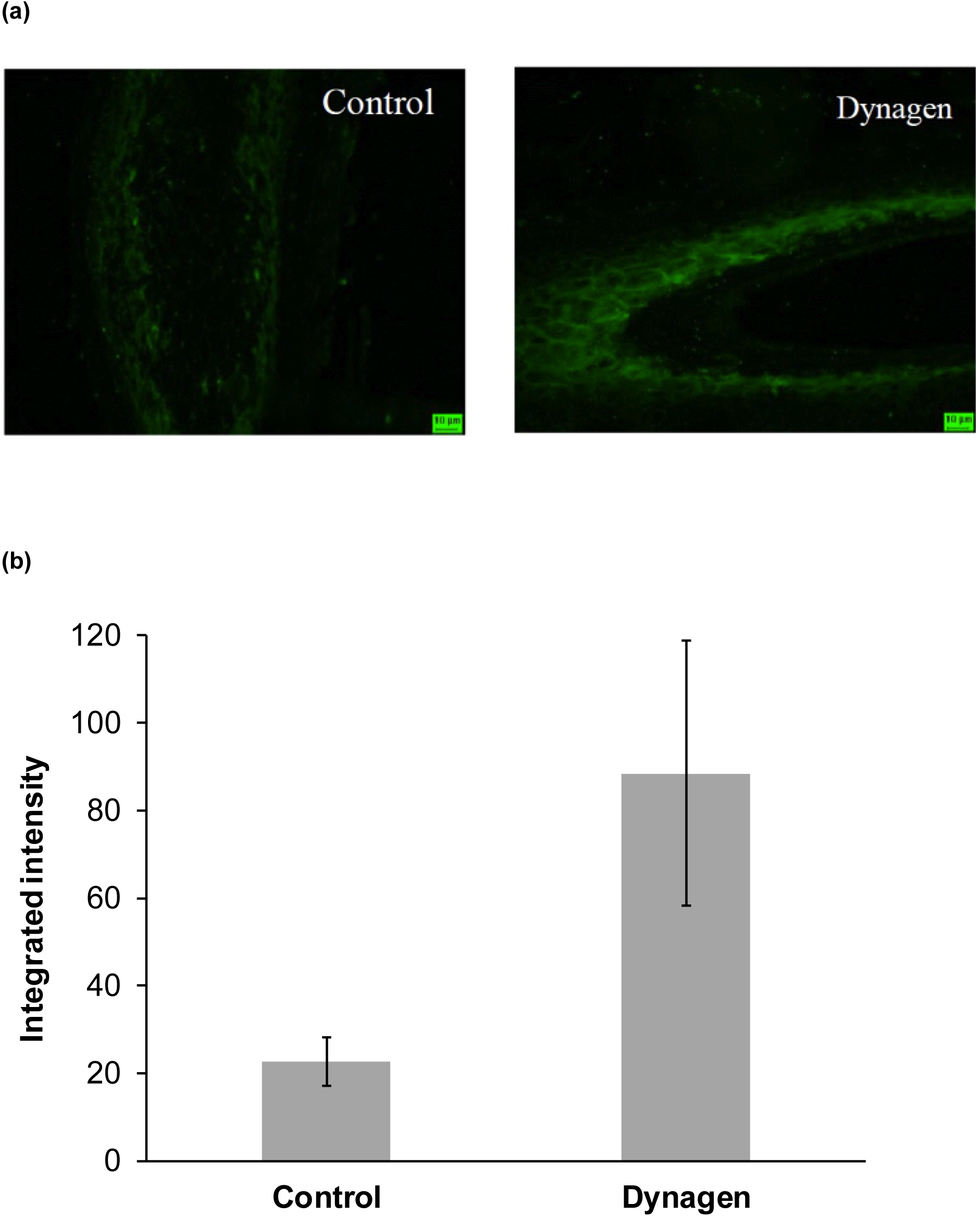
Claudin-1 expression in control and Dynagen-treated hair follicles cultured *in vitro*. (a) Representative photomicrographs illustrating claudin-1 localization. (b) Quantification of claudin-1 immunostaining. Data shown are means and standard errors of the mean for 180 regions analyzed in a total of nine follicles from three volunteers per group.

#### Hair-plucking force

Pig skin hair treated with the active LOT (n=254 hairs) required a significantly greater plucking force to remove hairs from the skin compared with that for the control-treated skin (n=251 hairs) (*P*<0.0001; Fig. 7).

**Figure 7.**
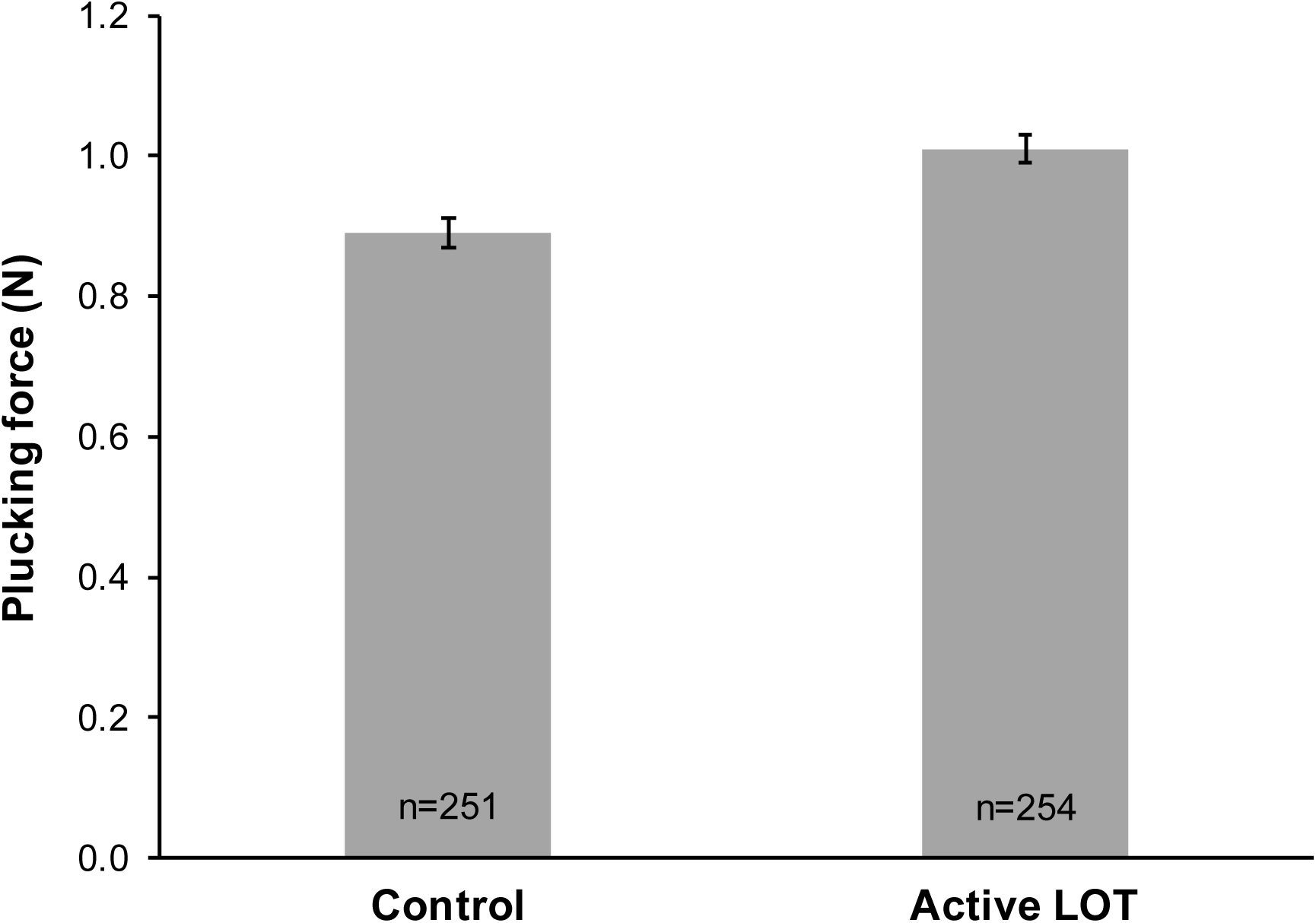
Effect of active LOT on the force required to pluck individual hairs from pig skin *ex vivo*. Data shown are means and standard errors of the mean. LOT = leave-on treatment; n = number of hairs.

#### Protease inhibition

The zinc salts included in the active LOT significantly reduced protease activity (*P*<0.05) compared with zinc salt-free control (Fig. 8).

**Figure 8.**
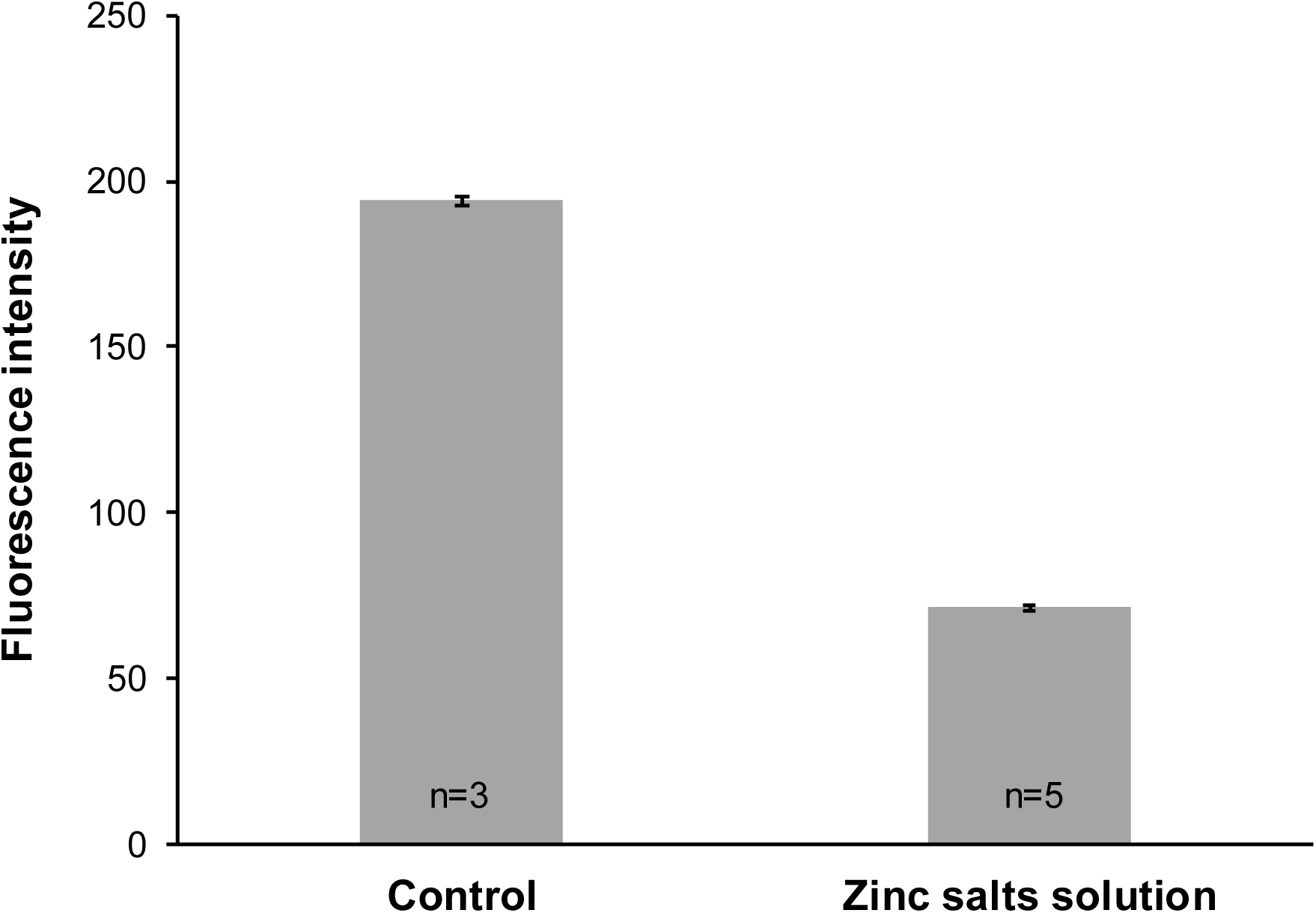
Protease inhibition with zinc salts solution compared with control. Intensity of fluorescence is proportional to protease concentration. Data shown are means and standard errors of the mean.

### Hair-fibre diameter

The mean diameter of single fibres (n=50) and the mean cross-sectional area of switches (n=5) treated with the active LOT increased by 2.49 µm (3.4%; *P*<0.0001) and by 0.14 mm^2^ (2.3%; *P*<0.0001), respectively, compared with the pre-treatment baseline (Table I). To establish whether this can be explained by the presence of glycerol in the Dynagen, the extent of fibre diameter increase associated with glycerol alone was determined using fresh single fibres (n=50) and switches (n=5). The mean fibre diameter and mean bundle thickness increased by 0.89 μm (1.2%; *P*=0.02) and by 0.03 mm^2^ (0.6%; *P*=0.004), respectively (Table I), after treatment with 0.45% glycerol solution, which is the effective glycerol concentration in the active LOT. Thus glycerol only accounts for ∼30% of the hair thickening seen with the active LOT.

**Table I.** Single hair-fibre diameter and hair-bundle thickness (cross-sectional area) following treatment with active LOT or glycerol

|  | Before treatment | After treatment | Change (after – before) | % increase | *P*-value<sup>a</sup> |
| --- | --- | --- | --- | --- | --- |
| **Mean fibre diameter (µm)** |  |  |  |  |  |
| Active LOT | 74.13 | 76.62 | 2.49 | 3.4 | <0.0001 |
| Glycerol (0.45%) | 75.69 | 76.58 | 0.89 | 1.2 | 0.02 |
| **Mean hair-bundle thickness (mm<sup>2</sup>)** |  |  |  |  |  |
| Active LOT | 6.00 | 6.14 | 0.14 | 2.3 | <0.0001 |
| Glycerol (0.45%) | 5.86 | 5.89 | 0.03 | 0.6 | 0.004 |

<sup>a</sup>One-tailed *t* test.
LOT = leave-on treatment; active LOT = LOT containing 1.5% Dynagen and zinc salts (0.0001% zinc gluconate, 0.018% zinc sulphate heptahydrate).

## Discussion

The studies reported in this paper indicate that an active LOT containing Dynagen plus zinc salts has beneficial effects on hair, including increasing hair-fibre diameter (as measured by laser-scanning micrometry), hair-follicle strength (as indicated by the tight-junction marker claudin-1 and the *ex vivo* hair-plucking force) and hair growth (as shown by immunohistochemical, *in vitro* and clinical studies).

### Hair growth

PTG assessments of hair growth rate in a cross-sectional analysis of 255 women confirmed that hair growth rate decreases with increasing age; however the data suggest that hair width increases with age to around age 40–45 years, before decreasing. This may offer an explanation for why no differences in average hair width were seen between the younger and older subject groups.

Comparison of the hair growth benefit associated with the active LOT (0.53 µm h^−1^) versus the age-related decrease in hair growth (0.043 µm h^−1^ per year) suggests that the use of Dynagen plus zinc salts can be translated into the equivalent of reducing the effects of this aspect of ageing by more than 12 years.

An effect on hair growth was also indicated by a significant increase in the expression of Ki67 in the hair bulbs obtained from subjects who had used the active LOT compared with those from subjects treated with control. This increase may be related to the increased hair growth as measured by PTG. The Ki67 protein is a reliable marker of cell proliferation, being present in all active phases of the cell cycle and absent in quiescent or resting phases [28–30]. Its expression is associated with increased hair growth, showing a decrease in Ki67-positive hair-matrix keratinocytes at the onset of catagen that has been linked to changes in differentiation markers in the outer and inner root sheaths [31], and increasing or decreasing in association with changes in hair-follicle proliferation in response to caffeine or testosterone, respectively [32, 33].

The mechanism of growth stimulation caused by the active LOT is unclear at present. It may be related to increased expression of proteins contributing to follicle structure (e.g. claudin-1) or to the addition of dipeptides as an extra nutrient source in the culture medium, supplying amino acids to the hair-follicle matrix, which then results in increased activity of these cells (as shown by the increase in Ki67 expression).

### Hair fall

Application of the zinc salts used in the active LOT significantly inhibited protease activity *in vitro*, and application of the active LOT increased the force required to pluck hair from pig skin; both findings indicate a more firmly anchored hair fibre. As the shear strain required to pluck hair from older subjects (aged >50 years) is significantly less than that required in younger subjects (both aged 20–50 years and <20 years) [12], strengthening the follicular root in this way provides an important benefit in the context of age-related changes to hair.

The well-known hair-growth cycle of anagen, catagen, telogen and exogen is accompanied by changes in the mechanical forces holding the hair shaft within the follicle. At the onset of telogen, the hair fibre is firmly held in the follicle. As this phase progresses the action of proteolytic enzymes results in a decrease in the proteins that serve to retain the hair fibre in place. The end result of this is that hair fibres are shed from the follicle due to the action of washing or brushing, as these forces exceed the forces holding the fibre in place. Inhibition of the hair follicle proteases delays exogen and results in retention of telogen fibres on the scalp. Measurement of the force required to pluck hair from a model skin provides an indication of the ability of a formulation to inhibit these enzymes and help retain hair fibres on the scalp [17, 18].

In the present study, hair-plucking force was measured on pig skin. One of the advantages of this is the availability of material from sources used for food production. However, some care has to be taken with data interpretation and extrapolation to human tissue. Pig skin is known to have a higher percentage of hairs in the telogen phase (∼50% telogen). This is an advantage from the point of view of examining the impact of protease inhibition, but may provide an overestimate of the impact that can be achieved in humans (∼10% hairs in telogen).

The junctional system in hair follicles is complex but may provide architectural stability to the skin, which is challenged mechanically and by ultraviolet light irradiation in daily life, and might also fulfil further functions, in particular contributing to the barrier properties of the skin [34]. The present studies have demonstrated that the active LOT increases the expression of claudin-1 (a key protein component of tight junctions). This, taken together with the increased expression of collagen IV and keratin 14 [24], is further evidence that the active LOT exerts its action via strengthening the structure of the hair follicle.

### Fibre diameter

The active LOT had a direct effect on the hair fibre, resulting in highly significant increases in diameter and hair-bundle cross-sectional area. It could be argued that both effects may be related to penetration of the dipeptides and glycerol from Dynagen into the fibre, resulting in swelling and physical coating of polymers from the formulation. However, the effect of the glycerol present in the Dynagen on swelling of the hair fibres was found to be substantially lower than that associated with the active LOT, only accounting for ∼30% of the hair thickening. It is possible that other ingredients in the formulation may contribute to the hair-thickening effect.

## Conclusions

Treatment with Dynagen plus zinc salts significantly increased single hair-fibre diameter and hair-bundle cross-sectional area. Hair-follicle integrity, as demonstrated by increased expression of the tight-junction marker claudin-1 and increased *ex vivo* hair-plucking force, was also increased following application of Dynagen. Application of active LOT (or Dynagen) was also associated with stimulation of hair growth, as indicated by results from tissue samples (Ki67 expression), cultured hair follicles and in the clinical study, in which the observed increased hair growth rate effect was equivalent to the reversal of 12 years of chronological ageing. Together, the results from this series of studies suggest that leave-on hair formulations containing Dynagen plus zinc salts provide the potential to ameliorate or mitigate age-related changes in hair, increasing hair growth and abundance, and follicular strength.

## Acknowledgments

Lynette Weddell was the Clinical Study Coordinator; Carl Donovan provided additional statistical analytical support; and Yunkun Xue conducted hair switch diameter measurements. Clinical studies were carried out at Alba Science Ltd, Edinburgh, UK

Editorial assistance was provided by Dr Duncan Porter of Anthemis Consulting Ltd, funded by Unilever R&D, Bebington, UK, in accordance with Good Publication Practice (GPP3) guidelines.

## Disclosure statement

All authors are employees of Unilever plc.

## Funding

The studies reported in this paper were funded by Unilever Research and Development.

## References

1. Sinclair, R. Diffuse hair loss. Int. J. Dermatol. 38 Suppl 1, 8–18 (1999).

2. Van Neste, D. Thickness, medullation and growth rate of female scalp hair are subject to significant variation according to pigmentation and scalp location during ageing. Eur. J. Dermatol. 14, 28–32 (2004).

3. Sinclair, R., Chapman, A. and Magee, J. The lack of significant changes in scalp hair follicle density with advancing age. Br. J. Dermatol. 152, 646–649 (2005).

4. Tajima, M., Hamada, C., Arai, T., Miyazawa, M., Shibata, R. and Ishino, A. Characteristic features of Japanese women’s hair with aging and with progressing hair loss. J. Dermatol. Sci. 45, 93–103 (2007).

5. Messenger, A.G. Hair through the female life cycle. Br. J. Dermatol. 165 Suppl 3, 2–6 (2011).

6. Robbins, C., Mirmirani, P., Messenger, A.G., Birch, M.P., Youngquist, R.S., Tamura, M., Filloon, T., Luo, F. and Dawson, T.L. Jr. What women want – quantifying the perception of hair amount: an analysis of hair diameter and density changes with age in Caucasian women. Br. J. Dermatol. 167, 324–332 (2012).

7. Kim, J.E., Lee, J.H., Choi, K.H., Lee, W.S., Choi, G.S., Kwon, O.S., Kim, M.B., Huh, C.H., Ihm, C.W., Kye, Y.C., Ro, B.I., Sim, W.Y., Kim, D.W., Kim, H.O. and Kang, H. Phototrichogram analysis of normal scalp hair characteristics with aging. Eur. J. Dermatol. 23, 849–856 (2013).

8. Kim, S.N., Lee, S.Y., Choi, M.H., Joo, K.M., Kim, S.H., Koh, J.S. and Park, W.S. Characteristic features of ageing in Korean women’s hair and scalp. Br. J. Dermatol. 168, 1215–1223 (2013).

9. Liu, C., Yang, J., Qu, L., Gu, M., Liu, Y., Gao, J., Collaudin, C. and Loussouarn, G. Changes in Chinese hair growth along a full year. Int. J. Cosmet. Sci. 36, 531–536 (2014).

10. Reddy, S.K. and Garza LA: The thinning top: why old people have less hair. J. Invest. Dermatol. 134, 2068–2069 (2014).

11. Tang, W., Zhang, S.G., Zhang, J.K., Chen, S., Zhu, H. and Ge, S.R. Ageing effects on the diameter, nanomechanical properties and tactile perception of human hair. Int. J. Cosmet. Sci. 38, 155–163 (2016).

12. El-Rifaie, A.A., Abdel Wehab, A.M., Gohary, Y. and El-Rifaie, A.E. The trichotillometry: a technique for hair assessment. Skin Res. Technol. 22, 15–19 (2016).

13. Ellis, J.A. and Sinclair, R.D. Male pattern baldness: current treatments, future prospects. Drug Discov. Today 13, 791–797 (2008).

14. Gubelin Harcha, W., Barboza Martinez, J., Tsai, T.F., Katsuoka, K., Kawashima, M., Tsuboi, R., Barnes, A., Ferron-Brady, G. and Chetty, D. A randomized, active- and placebo-controlled study of the efficacy and safety of different doses of dutasteride versus placebo and finasteride in the treatment of male subjects with androgenetic alopecia. J. Am. Acad. Dermatol. 70, 489–498.e483 (2014).

15. Kelly, Y., Blanco, A. and Tosti, A. Androgenetic alopecia: an update of treatment options. Drugs 76, 1349–1364 (2016).

16. Takahashi, T., Kamimura, A., Yokoo, Y., Honda, S. and Watanabe, Y. The first clinical trial of topical application of procyanidin B-2 to investigate its potential as a hair growing agent. Phytother. Res. 15, 331–336 (2001).

17. Higgins, C.A., Westgate, G.E. and Jahoda, C.A. Modulation in proteolytic activity is identified as a hallmark of exogen by transcriptional profiling of hair follicles. J. Invest. Dermatol. 131, 2349–2357 (2011).

18. Bhogal, R.K., Mouser, P.E., Higgins, C.A. and Turner, G.A. Protease activity, localization and inhibition in the human hair follicle. Int. J. Cosmet. Sci. 36, 46–53 (2014).

19. Debela, M., Goettig, P., Magdolen, V., Huber, R., Schechter, N.M. and Bode, W. Structural basis of the zinc inhibition of human tissue kallikrein 5. J. Mol. Biol. 373, 1017–1031 (2007).

20. Westgate, G.E., Shaw, D.A., Harrap, G.J. and Couchman, J.R. Immunohistochemical localization of basement membrane components during hair follicle morphogenesis. J. Invest. Dermatol. 82, 259–264 (1984).

21. Couchman, J.R. and Gibson, W.T. Expression of basement membrane components through morphological changes in the hair growth cycle. Dev. Biol. 108, 290–298 (1985).

22. Messenger, A.G., Elliott, K., Temple, A. and Randall, V.A. Expression of basement membrane proteins and interstitial collagens in dermal papillae of human hair follicles. J. Invest. Dermatol. 96, 93–97 (1991).

23. Langbein, L., Yoshida, H., Praetzel-Wunder, S., Parry, D.A. and Schweizer, J. The keratins of the human beard hair medulla: the riddle in the middle. J. Invest. Dermatol. 130, 55–73 (2010).

24. Dal Farra, C., Domloge, N. and Botto, J.-M. Use of yeast peptide hydrolysate as an active agent for strengthening hair. United States patent application No. US 2012/0220541 A1. United States Patent and Trademark Office (2012). Available from: http://pdfaiw.uspto.gov/.aiw?PageNum=0&docid=20120220541; accessed March 2017.

25. Canfield, D. Photographic documentation of hair growth in androgenetic alopecia. Dermatol. Clin. 14, 713–721 (1996).

26. Philpott, M.P., Green, M.R. and Kealey, T. Human hair growth *in vitro*. J. Cell Sci. 97, 463–471 (1990).

27. Cohen, B. The cross-section trichometer: a new device for measuring hair quantity, hair loss, and hair growth. Dermatol. Surg. 34, 900–910 (2008).

28. Gerdes, J., Schwab, U., Lemke, H. and Stein, H. Production of a mouse monoclonal antibody reactive with a human nuclear antigen associated with cell proliferation. Int. J. Cancer. 31, 13–20 (1983).

29. Gerdes, J., Lemke, H., Baisch, H., Wacker, H.H., Schwab, U. and Stein, H. Cell cycle analysis of a cell proliferation-associated human nuclear antigen defined by the monoclonal antibody Ki-67. J. Immunol. 133, 1710–1715 (1984).

30. Scholzen, T. and Gerdes, J. The Ki-67 protein: from the known and the unknown. J. Cell Physiol. 182, 311–322 (2000).

31. Commo, S. and Bernard, B.A. Immunohistochemical analysis of tissue remodelling during the anagen-catagen transition of the human hair follicle. Br. J. Dermatol. 137, 31–38 (1997).

32. Fischer, T.W., Hipler, U.C. and Elsner, P. Effect of caffeine and testosterone on the proliferation of human hair follicles *in vitro*. Int. J. Dermatol. 46, 27–35 (2007).

33. Fischer, T.W., Herczeg-Lisztes, E., Funk, W., Zillikens, D., Biro, T. and Paus, R. Differential effects of caffeine on hair shaft elongation, matrix and outer root sheath keratinocyte proliferation, and transforming growth factor-beta2/insulin-like growth factor-1-mediated regulation of the hair cycle in male and female human hair follicles *in vitro*. Br. J. Dermatol. 171, 1031–1043 (2014).

34. Brandner, J.M., McIntyre, M., Kief, S., Wladykowski, E. and Moll, I. Expression and localization of tight junction-associated proteins in human hair follicles. Arch. Dermatol. Res. 295, 211–221 (2003).

